# AFCMEasyModel: An Easy Interface for Modeling Competing Endogenous RNA Networks using ODEs

**DOI:** 10.1101/241026

**Authors:** Mohammad M. Tarek

## Abstract

Competing endogenous RNA networks have been considered to be important regulators of genetic data expression. Circular RNAs and microRNAs interact to form a circular sponge that have been shown to regulate messenger RNAs and hence regulating gene expression. The kinetics by which these non-coding RNAs interact together affecting gene expression are crucial to understand the mechanism of their regulatory function. Herein, we developed AFCMEasyModel as a user-friendly shiny app that enables users to modify regulation parameters of a competing endogenous RNA network based on interaction between circular RNAs and microRNAs in the simulation environment to form a sponge complex. The App provides the source-code for more customized models and allow users to download simulation plots for supplementation of their publications.

The App was made available for public-access at: https://mohammadtarek.shinyapps.io/afcmeasymodel/

## Introduction

Competing endogenous RNA (ceRNA) has been recognized as one of the major regulators of genetic expression.^[1]^ CeRNAs play significant rules through indirect interactions among RNA transcripts Circular RNAs (circRNAs) are non-coding RNAs class (ncRNAs) that has shown strong evidence for gene expression regulation due to the growing research data about it is interaction in transcriptional and posttranscriptional epigenetic regulation. circRNAs have been proved to perform their regulatory function in a competing endogenous RNA (ceRNA) fashion.^[2,3]^ CircRNAs model as ceRNAs indicated a strong role for regulating miRNAs activity which has been found to be associated different diseases.^[4,5]^ MiRNAs have been known to participate in post transcriptional modifications of target mRNAs through an RNA-induced silencing complex (RISC) in which miRNAs long seed regions bind miRNA regulatory elements MREs in the 3'UTR.^[5,6]^ The complexity of ceRNAs interactions has been considered a very challenging molecular relationship to understand because of variable dependency on multi factors regarding the interaction complex. These multifactorial-dependent interaction may be affected by the degree of complementarity between MREs and long seed regions as well as the degree of interaction between Argonaute proteins to miRNAs that could stimulate functional segments on miRNA transcripts which could later affect the downregulation effect on target mRNAs.^[7]^ The complexity of RNA regulation networks has been observed to be indicative for miRNA mediated silencing for a specific target mRNA where the concentration of that miRNA tends to be target-focused for a specific misregulation of single target from the set of miRNA targets.^[8–10]^ Although miRNAs could potentially act as a limiting factor for RNA regulatory Networks, Literature also found a strong evidence of network regulatory effect where miRNAs show indirect interactions with a set of mRNAs.^[11]^ These networks have further elaborated the crosstalk effect among molecular RNA species, however, these regulatory interactions are still dependent on the MREs shared by target mRNAs in a network.^[12,13]^ The nature of RNA crosstalks opens the door to a molecular environment that is very susceptible to stochastic effects, in which mathematical models could give us detailed insights about the RNA system kinetics that could be further translated into therapeutic intervention studies to regain genetic equilibrium into these regulatory networks. CeRNA regulatory networks have been observed to be of significant carcinogenic implications in different human tumors.^[14,15]^ While circRNAs perform their function in a sponge like manner, they also acquired some interesting functional properties such as tissue specificity, stability and abundance in clinical diagnostic samples. R language has been enriched with different packages that facilitated the simulation of biomedical data making good use of current computational processing abilities.^[16]^ Herein, we aimed to develop an interactive plotting app for simulating ceRNA networks that includes an interacting molecular miRNA specie, an mRNA specie as well as a circular RNA specie. It would be also valuable for researchers to have an interactive framework to test ceRNA regulatory simulations in real time.

## Implementation

The tool was implemented in R language using shiny platform.^[17]^ The app depends on solving ordinary differential equations ODEs that integrates all molecular species with corresponding kinetics parameters that were obtained from literature [Table-1].^[18]^ DeSolve R package has been used as the main solver of a customized system of ODEs [Figure-1].^[19]^ A custom function called “IGEM” was created that includes three ODEs in which each equation corresponds to the kinetics of distinct molecular specie in the simulation environment of ceRNA network.

**Figure-1.**
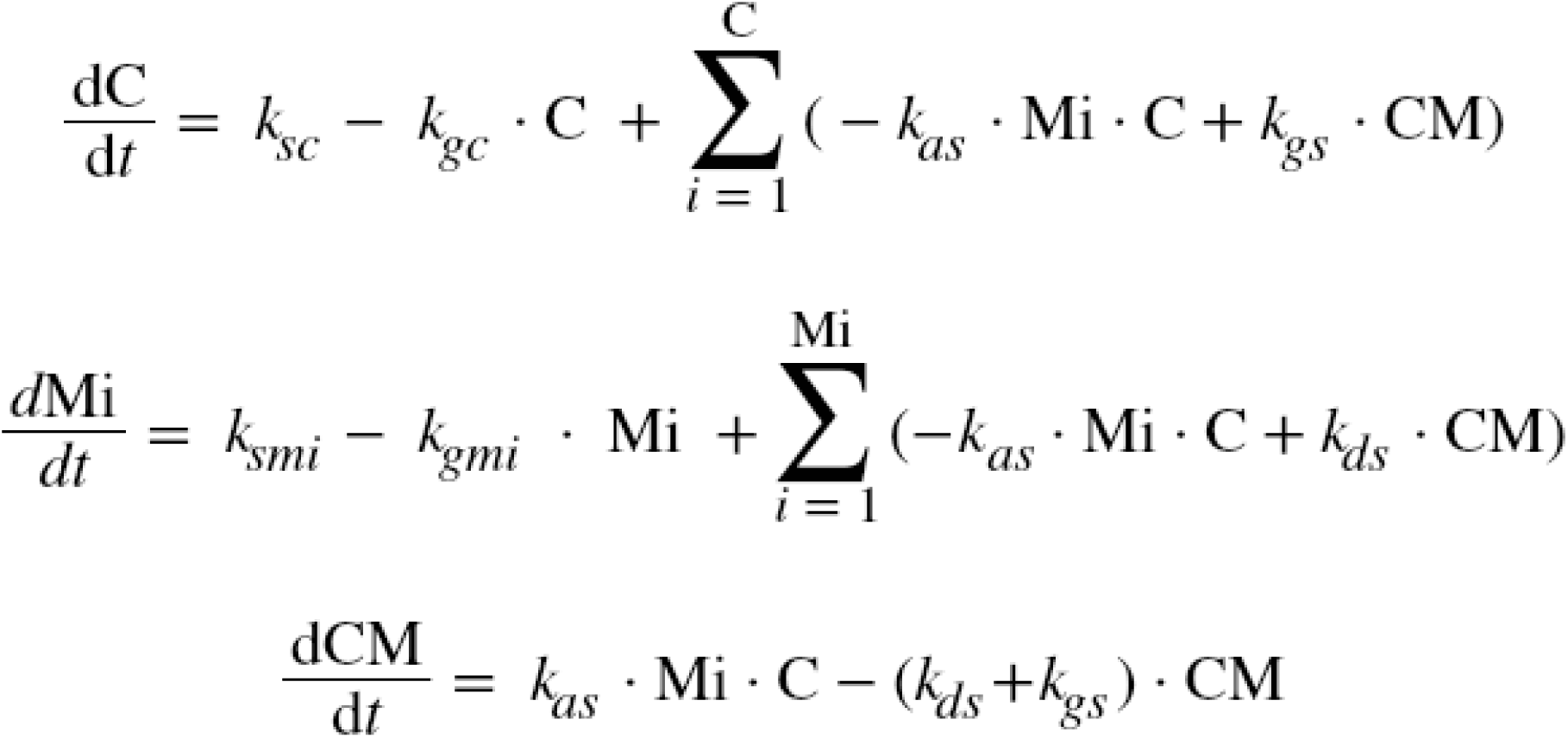
Set of Ordinary Differential Equations Describing the system

**Table-1.**
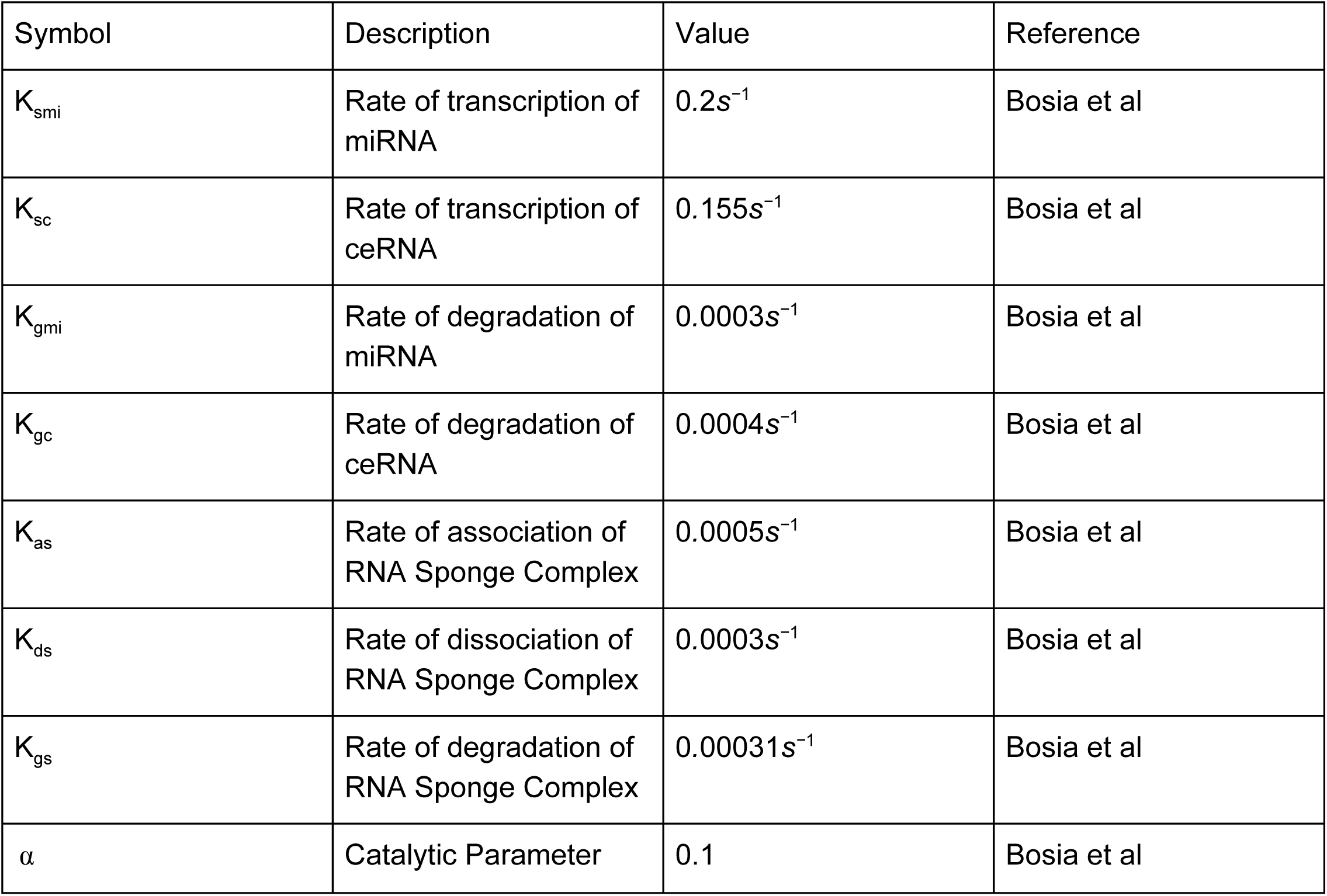
Parameter values of ceRNA network regulation

## Design

The web tool was designed according to the user interface previously developed at AFCMHeatMap tool.^[20]^ On the left, users can modify different regulation parameters such as Initial concentrations of each molecular specie (circRNA, miRNA and ceRNA sponge), rates of synthesis, degradation rates and simulation time. The Simulation plot frame was designed to be occupying the whole main panel for better visualization. The R package ggplot2^[21]^ was used with reshape function to adjust the data frame representing the numerical solving of ODEs in the simulation plot.

## Input

The slider input makes it very simple for users to modify these parameters in real time when the plot is updated on the main panel to the right for each slight modification the users perform to the slider controls on the left. The source-code provided on the application github-repository provides users with the ability to create new functions with modified systems of ODEs to fit in different simulation parameters or simulate a different regulation model. For users to resimulate another model, they would just modify the the slider controls for an updated plot. Users also may just refresh the browser page for another new simulation session.

## Output

The simulation plot [Figure-2] appears in a grey grid background where the x-axis represents simulation time, while the y-axis represents the concentration of molecular species of the ceRNA network over-time. The red line represents change in circRNA concentration per time, the green line represents the change in miRNA concentration in the simulation environment per time while the blue line represents the change in sponge complex concentration per time. The user then could easily download the simulation plot by saving it as image as it is compatible to *.PNG extension

**Figure-2.**
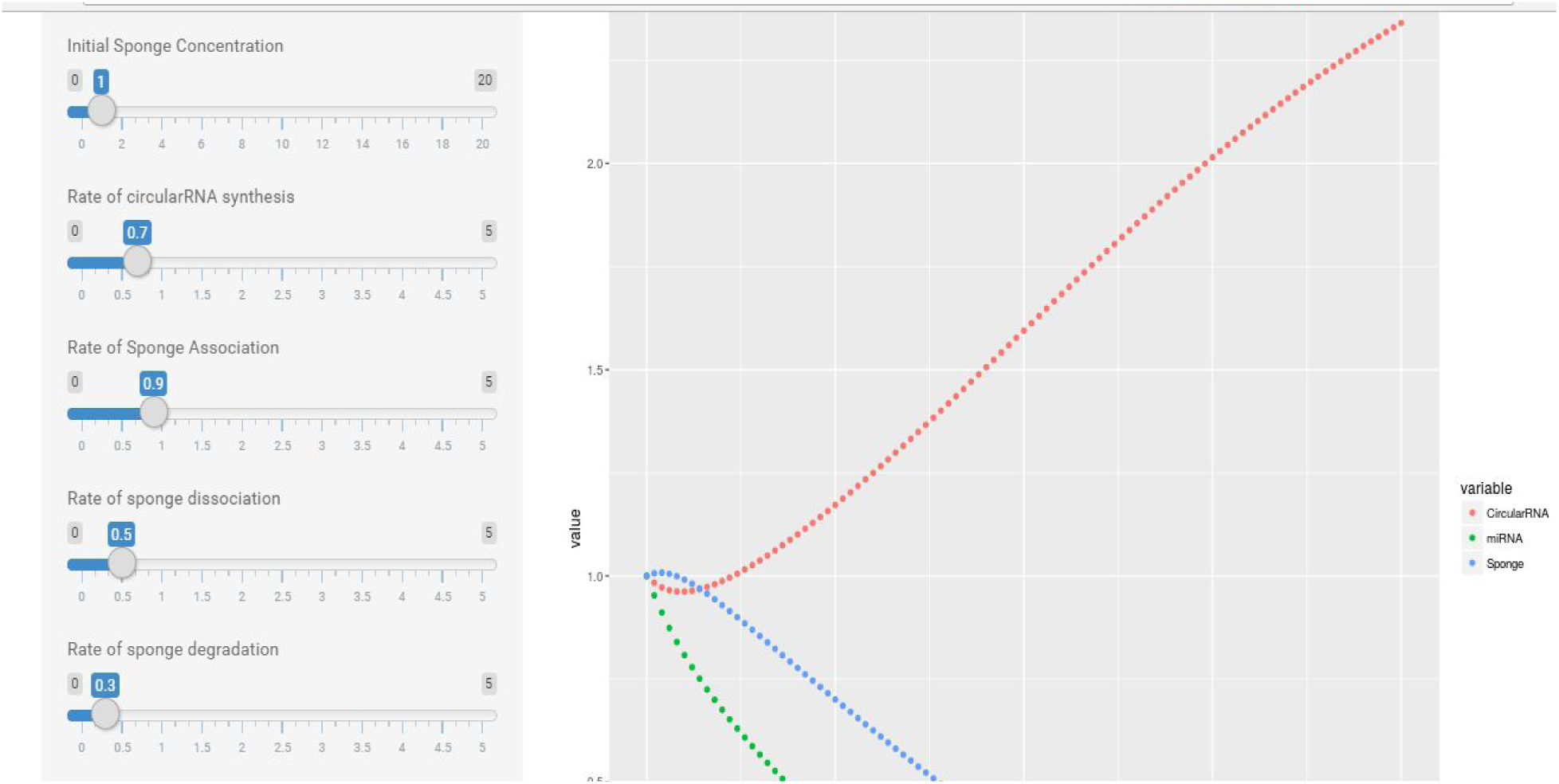
A user interactive model generated y AFCMEasyModel

## User-Developed Model

This Model [Figure-3] aims to describe the regulation of competing endogenous RNA (ceRNA) network using ordinary differential equations to get insights about the kinetics of molecular species inside the network. The Model was built on Synthetic biology markup language SBML standards to describe biological parts and their interactions including: transcription, degradation, association and dissociation of both the ceRNA and miRNA. The Model describes an inhibitory relationship, where the miRNA binding to ceRNA inhibits the miRNA action on its target mRNA. We can estimate that effect from the change in free miRNAs in the simulation run. This model was constructed as a part of AFCM-Egypt team project modelling that participated at IGEM (International Genetically Engineered Machine Competition) in 2017.

**Figure-3.**
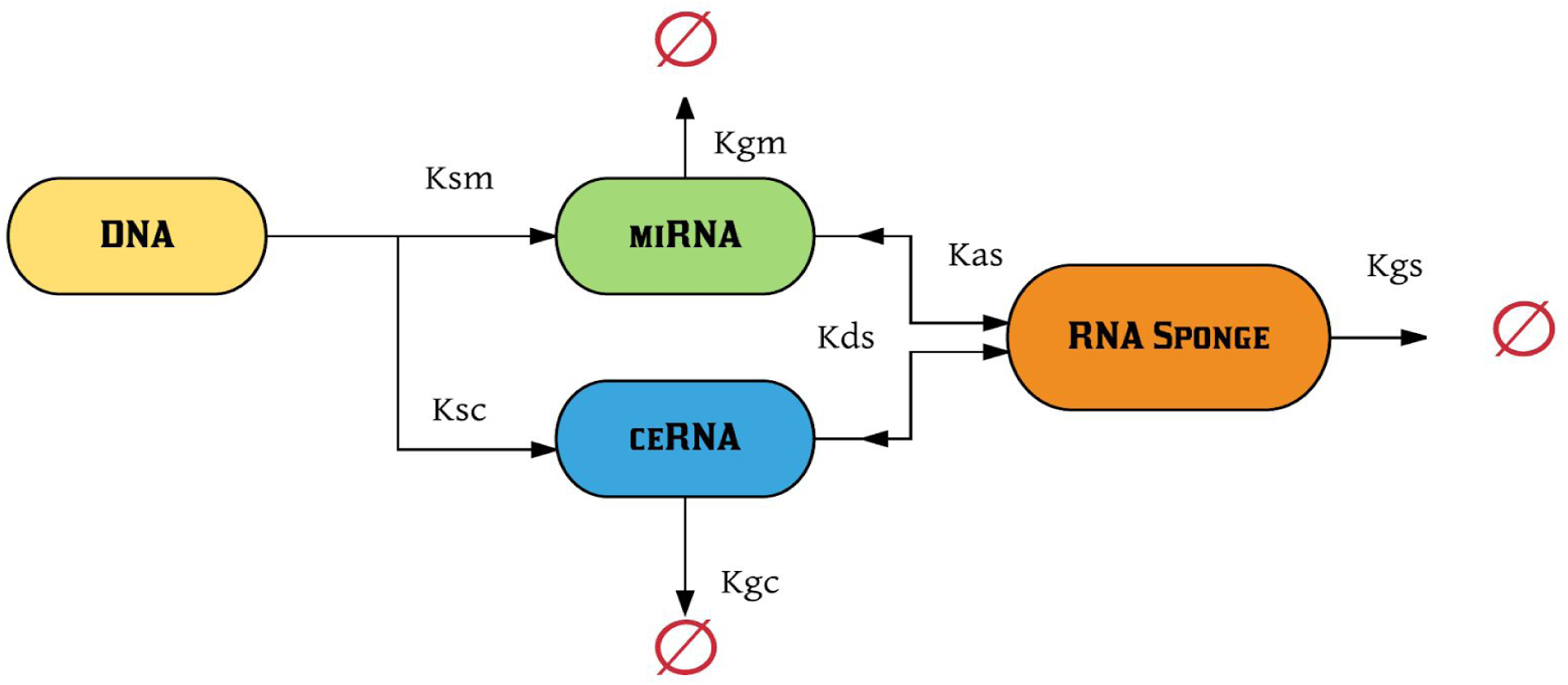
Graphical representation for ceRNA model

## Limitations and Future Perspective

As previously mentioned in the *Introduction*, miRNAs potentiate extensive crosstalk effects among RNA regulatory network. Due to the low concentration of these RNA molecular species in the simulation environment, stochasticity could play an important role in the interaction nature which could lead to further phenotypic variation on the results of gene expression. Therefore, we intend to add another feature of stochastic simulation based on Gillespie algorithm to AFCMEasyModel to help modeling stochastic effects. Another considerable Limitation of this tool as we see is the modeling time, which obviously could be manipulated through debugging the original code on R platform depending on the user processing abilities rather than a limited processing server.

## Conclusions

The behavioral changes of ceRNA networks should consider the different mechanisms by which ncRNAs - such as miRNAs or circRNAs - could act in their nuclear or cytosolic environments to achieve their regulatory function. The simple system of ODEs that was implemented in AFCMEasyModel focused on temporal reciprocity character of miRNAs action on target mRNAs. The circular sponge formation described the functional regulation of miRNA that could be formed after association with circular RNA, The app considered degradation of each molecular species as well as degradation of sponge complex. The catalytic activity in the reaction environment was represented by user modifications to the synthesis rates in the limited time frame of 100 time unit which could be modified by users through manipulating the source-code provided in form of server R file and user-interface R file on an external R compiler. Therefore, AFCMEasyModel provided a simple and interactive simulator for ceRNA networks of circRNAs and miRNAs that could be further developed to include parametric representations of ncRNAs crosstalks as well as stochastic effects that could be induced by these crosstalks in the simulation environment.

## Availability and requirements

- **App name:** AFCMEasyModel
- **App home page:** https://mohammadtarek.shinyapps.io/afcmeasymodel/**Requires**: Only requires an Internet Connection.
- **Programming language:** R Shiny Framework
- **License:** GNU General Public License version 3.0 (GPL-3.0)

### Availability of Source-Code files

All source-code files were made available on Github at: https://github.com/MTarekM/AFCMeasymodel

